# Attrition Rate in Infant fNIRS Research: A Meta-Analysis

**DOI:** 10.1101/2021.06.15.448526

**Authors:** Sori Baek, Sabrina Marques, Kennedy Casey, Meghan Testerman, Felicia McGill, Lauren Emberson

## Abstract

Understanding the trends and predictors of attrition rate, or the proportion of collected data that is excluded from the final analyses, is important for accurate research planning, assessing data integrity, and ensuring generalizability. In this pre-registered meta-analysis, we reviewed 182 publications in infant (0-24 months) functional near-infrared spectroscopy (fNIRS) research published from 1998 to April 9, 2020 and investigated the trends and predictors of attrition. The average attrition rate was 34.23% among 272 experiments across all 182 publications. Among a subset of 136 experiments which reported the specific reasons of subject exclusion, 21.50% of the attrition were infant-driven while 14.21% were signal-driven. Subject characteristics (e.g., age) and study design (e.g., fNIRS cap configuration, block/trial design, and stimulus type) predicted the total and subject-driven attrition rates, suggesting that modifying the recruitment pool or the study design can meaningfully reduce the attrition rate in infant fNIRS research. Based on the findings, we established guidelines on reporting the attrition rate for scientific transparency and made recommendations to minimize the attrition rates. We also launched an attrition rate calculator (**LINK**) to aid with research planning. This research can facilitate developmental cognitive neuroscientists in their quest toward increasingly rigorous and representative research.

**Highlights:**

- Average attrition rate in infant fNIRS research is 34.23%
- 21.50% of the attrition are infant-driven (e.g., inattentiveness) while 14.21% are signal-driven (e.g., poor optical contact)
- Subject characteristics (e.g., age) and study design (e.g., fNIRS cap configuration, block/trial design, and stimulus type) predict the total and infant-driven attrition rates
- Modifying the recruitment pool or the study design can meaningfully reduce the attrition rate in infant fNIRS research

## 1 Introduction

### 1.1 Importance of attrition rate in fNIRS research

The introduction of fNIRS as a neuroimaging technique has transformed the landscape for infant brain research. fNIRS is an increasingly popular tool that non-invasively measures cortical hemodynamic responses (Elwell & Cooper, 2011). fNIRS is safe, affordable, portable, and increases infants’ task compliance compared to other neuroimaging techniques (for reviews see Aslin & Mehler, 2005; Lloyd-Fox et al., 2010; Gervain et al., 2011). A rapid development of analyses techniques (e.g., Behrendtet al., 2018) has helped fNIRS become relatively motion-robust (Lloyd-Fox et al., 2010; Nishiyori, 2016). Likely because of these advantages, a recent meta-analysis of infant neuroimaging showed that fNIRS studies were rapidly increasing in number even compared to other neuroimaging modalities (Azhari et al. 2020).

One major limitation of infant fNIRS research, however, is the attrition rate. Thoroughly understanding attrition is essential for the scientific integrity of the field. Attrition rate is defined as the proportion of collected data that was excluded from final analyses. High attrition rate is a prevalent issue in infant research regardless of methodology (e.g., Leroy et al., 2011; Picton and Taylor, 2007; Dehaene-Lambertz et al., 2010) and presents major challenges: A high attrition rate can lead to a smaller representative sample which threatens the external validity and reproducibility of results. High attrition also increases the cost of research. Understanding the attrition rate for fNIRS research and the factors that affect it is crucial to efficiently design scientific research, ensure external validity of results (Flick, 1988), and mitigate factors that increase attrition.

### 1.2 Possible causes of attrition in infant fNIRS research

#### 1.2.1 Infant-driven attrition

Infants’ shorter attention span and likeliness to become fidgety and agitated (e.g., Aylward et al., 2002; Borgaro et al., 2003; Marshall et al., 2009) may increase the attrition rate in infant research. Researchers attempt to reduce the attrition rate by limiting the number of experimental conditions and the duration of stimuli (e.g., Hoehl and Wahl, 2012) and using multimodal stimuli to make the stimuli more attention-grabbing (e.g., Richards, 2003; Reynolds & Guy, 2012).

However, evidence for the effectiveness of focusing on these factors to reduce attrition in infant research is mixed according to two previous meta-analyses. On one hand, Slaughter and Suddendorf (2007) found that the most common reason for attrition in infant behavioral studies (habituation-based looking time studies) is infant fussiness (13.7%, total attrition rate: 22.6%). This meta-analysis found no effect on attrition rate as a result of experimental design or subject age. On the other hand, Stets et al. (2012) analyzed attrition rate in infant EEG research and

found that the average attrition rate in infant EEG research was 47.3%. Unintuitively, they found that the overall attrition rate was 18.6% lower when auditory-only stimuli were used, instead of visual or audiovisual stimuli. This result is in direct contrast to the assumption in infant research that using multimodal stimuli can decrease attrition (e.g., Richards, 2003; Reynolds & Guy, 2012). These mixed conclusions from meta-analyses and common practice in the field emphasizes the need for systematic research on the factors that affect attrition.

#### 1.2.2 Signal-driven attrition

Cristia et al. (2013) conducted a review of the infant fNIRS literature and noted that fNIRS-specific factors affected the attrition rate. Studies using headsets with more than 20 optodes experienced very high attrition rates (upwards of 50%), although this finding may be driven by the small number of labs that used a large number of sensors in the early days of fNIRS. Also, fNIRS research on newborns and 5-to-8 month old infants experienced lower attrition rates compared to research on 2-to-4 month old infants, but it is not clear whether this finding is a result of infant-specific factors (e.g., fussiness) or fNIRS factors (e.g., hair, strength). This finding differs from those found in the two previously mentioned meta-analyses on attrition (Stets et al., 2012; Slaughter & Suddendorf, 2007) that found no effect of age.

### 1.3 Current meta-analysis

The current meta-analysis updates and extends these meta-analyses and reviews with a specific focus on fNIRS and attrition. We investigated an expanded number of potential predictors of attrition compared to previous meta-analysis and reviews. We examined whether major findings from Cristia et al (2013) replicate (e.g., age related effects) and include 7 more years of data, which represents a high proportion of the total publications in this young field. In addition, we investigated three types of attrition rates that could arise in a research setting: **Total Attrition Rate** (TAR; proportion of subjects who were excluded for any reason), **Infant-Driven Attrition Rate** (IAR; proportion of subjects who were excluded for the infant subjects’ behavior, such as fussiness), and **Signal-Driven Attrition Rate** (SAR; proportion of subjects who were excluded due to fNIRS signal-related issues such as poor optical contact). Although there may be other types of attrition rates that are also at play, we focused on these three types because they were the largest share of the attrition, were commonly disclosed, and were targets for improvement for future research.

We examined attrition rates and their predictors in three sections. First, we explored the overall trends in fNIRS research over the years reviewed. Second, we explored the possible effects of subject-related parameters on attrition rate (e.g., age). Third, we explored the effects of experimental design-related parameters (e.g., stimulus type). Then, we concluded a set of guidelines on how to report attrition rate to ensure transparent scientific results and suggestions how to best minimize the attrition rate to ensure replicable results with minimal data loss. We also have a visualization tool that is open to researchers who want to quantify estimated attrition rates for their studies based on our meta-analysis, find studies with the same experimental parameters and to consider how your experimental choices might influence your attrition rate. Together, this meta-analysis is aimed at increasing the validity and transparency of infancy and fNIRS research.

## 2 Material and Methods

### 2.1 Open-science

Prior to any of the following analyses, the scope, hypotheses, and analyses of this meta-analysis were pre-registered on Open Science Framework (OSF) Registries: http://osf.io/6a2vq. The entirety of this dataset can be accessed via https://github.com/soribaek/Attrition-Rate-in-Infant-fNIRS-Research. We have also launched an attrition rate calculator (**LINK**) to help estimate the attrition rate for an infant fNIRS experiment *a priori* to aid with research planning.

### 2.2 Literature search

To ensure a thorough and complete review of the literature, data collection for this meta-analysis used the methodology presented in “Searching for studies: A guide to information retrieval for Campbell Systematic Reviews” for comprehensive, transparent, reproducible, and unbiased data collection (Kugley et al., 2016). We designed the overall search strategy, parameters, language, validated the search, conducted and documented all searches, and deduplicated the search results in consultation with an experienced behavioral sciences librarian.

Database searching and snowballing provided the initial pool of studies to be screened for inclusion. We identified 10 studies that fully met the inclusion criteria to use as a validation set. The validation set was then used to design and validate custom searches in each of the following databases: Scopus, APA PsycINFO, and PubMed. A complete list of validation articles can be found in Appendix A. From this set of studies, the following search terms were identified from the title, abstracts, and keywords: fnirs, functional near infrared spectroscopy, nirs, near infrared spectroscopy, baby, babies, infant, infants, toddler. The search terms were connected with OR or AND, truncation devices were used where appropriate, and syntax was adjusted for each database. When available, database limiters were applied to limit the results to language (English) and document type (articles, conference papers, dissertations, and theses). There was no limitation placed on publication date. These data searches were conducted on April 3, 2020.

Searches were conducted, documented, and can be found in their entirety in Appendix B. A total of 3,812 records were identified via database searching and three records were obtained by requesting for data in ongoing research. 1,698 duplicates were removed. Thus, a total of 2,117 records were screened for inclusion. From the 2,117 records, authors screened the titles and abstracts to determine whether each record meets the following criteria: 1) human infant subjects, between 0 and 2 years old; 2) cognitive neuroscience research (i.e., looking at blood oxygenation as a proxy for neural activity), rather than a clinical or medical study (e.g. looking at oxygenation levels unrelated to neural activity); 3) empirical study, rather than reviews or case reports; 4) full-text available, rather than only abstracts; 5) presentation of new data, rather than a re-analysis of a previously published dataset. We excluded 1,901 hits using these criteria to result in a list of 216 publications for analysis.

Finally, after reading the 216 publications, we excluded 34 publications that did not report total or final sample size. Thus, we conducted the final analyses on 182 publications. The study selection process is articulated in Figure A. In these 182 publications, there were 272 experiments with distinct infant subject samples. Thus, on average, there were approximately 1.42 (range: 1 to 4) experiments per publication.

**Figure A.**
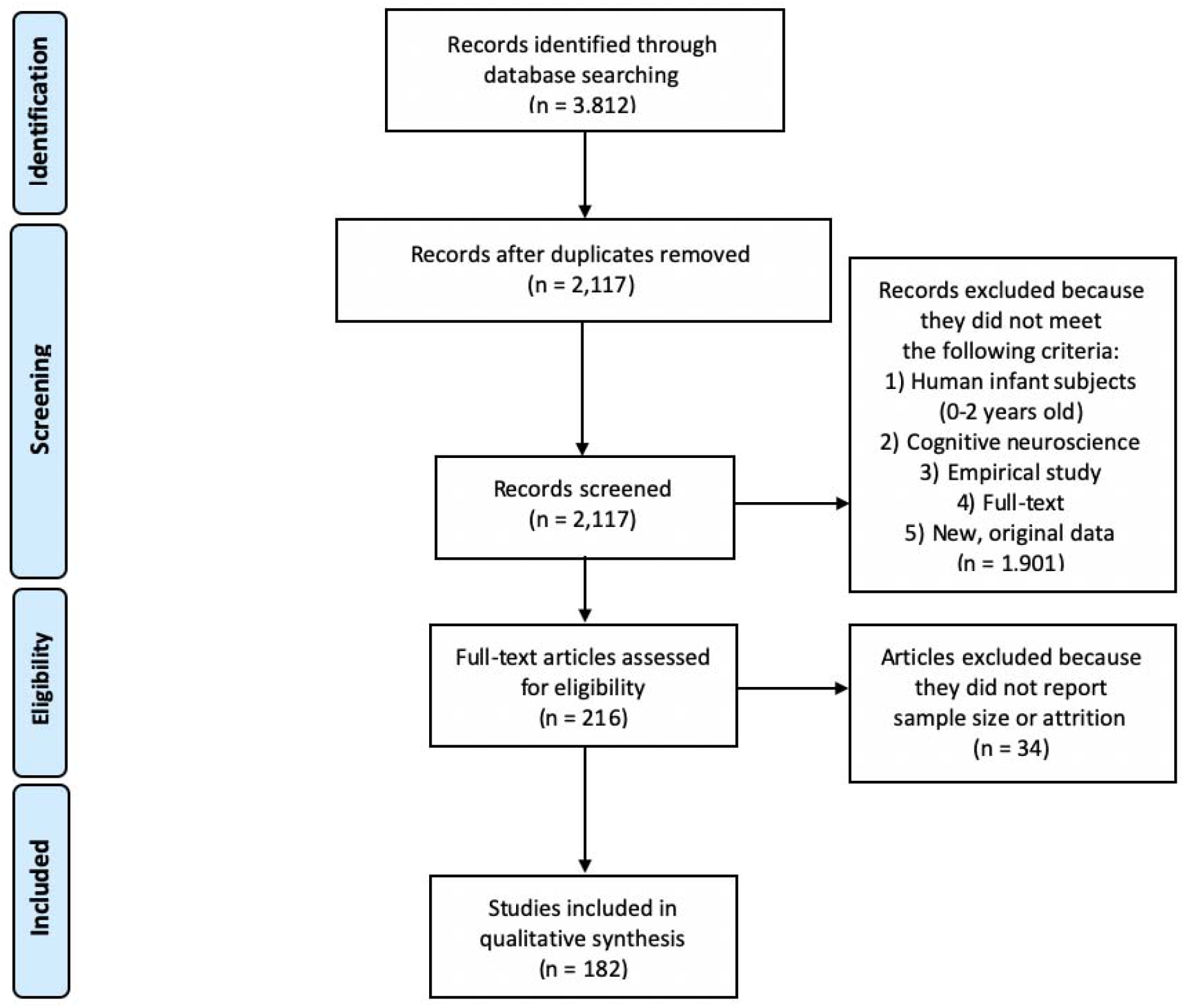
Study selection process. The study selection process is outlined here using the PRIMA Flow Diagram.

### 2.3 Attrition Rates

We defined the three types of attrition rates as follows to account for different types of data exclusion that could arise in a research setting. We used these definitions to calculate the attrition in each of the 272 experiments.

#### Total Attrition Rate (TAR)

Total Attrition Rate (TAR) referred to the total proportion of subjects who were excluded for any reason from the total recruitment pool. This was defined as the ratio of the number of subjects who remained in the final analysis sample over the total number of subjects who were recruited.

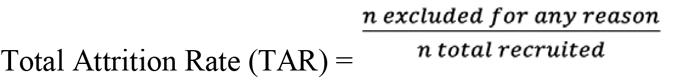

#### Infant-Driven Attrition Rate (IAR)

Infant-Driven Attrition Rate (IAR) referred to the proportion of subjects who were excluded for the infant subjects’ behavioral reasons, such as the subject resisting the fNIRS cap or being fussy and inattentive during the task.

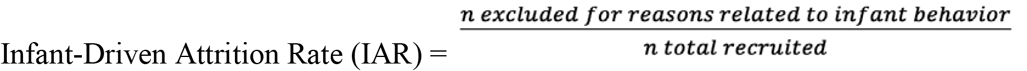

#### Signal-Driven Attrition Rate (SAR)

Signal-Driven Attrition Rate (SAR) referred to the proportion of subjects who were excluded for signal-related reasons, such as weak signals or strong artifacts.

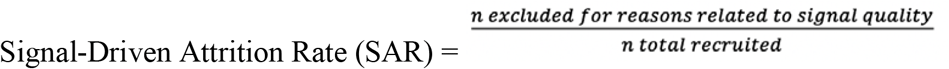

### 2.4 Parameters

#### 2.4.1 Subject Parameters

We defined Subject Parameters as characteristics of the infant subjects that may have influenced the attrition rate. We coded five Subject Parameters: **Age, sex, hospitalization status, preterm status**, and **clinical status**. Age was defined as the mean age in months of all infant subjects in the experiment. Sex was defined as the percentage of infants that were female among the subjects; although researchers in the field of developmental cognitive neuroscience are working to allow non-binary reporting of sex in future studies, our analyses were performed on publications that report only binary sex and, thus, we summarize sex in this way. Hospitalization status was binarily defined based on whether the study was conducted in the hospital (1) or not (0). Preterm status was defined based on whether the recruited infant subjects of each study were preterm (1) or full-term (0) infants, as determined by each experimenter’s criteria. Finally, the clinical status of the infants was coded as typically developing (0), at-risk, such as preterm or familial risk for autism spectrum disorder (1), or atypically developing, such as Down’s syndrome (2). All parameters that did not correspond to exactly one of these options were coded as “other” (−1); for example, some studies (Edwards et. al, 2017; Lloyd-Fox et. al, 2013) reported results from all subjects without differentiating among at-risk and low-risk subjects when reporting reasons for attrition, so this variable was coded as -1. There were 8 of these -1 values in the total dataset. They were removed from further analyses for clearer interpretability.

#### 2.4.2 Design Parameters

We defined *Design Parameters* as characteristics of each experimental study design that may have influenced the attrition rate. We coded for six Design Parameters: **fNIRS cap configuration, study design (i.e**., **whether blocks or trials were used), number of trials/blocks, length of trials/blocks, presence of stimuli**, and **type of stimuli**. fNIRS configuration was a continuous variable, defined as the number of channels used to collect data in each study. Study design was a binary variable, defined as trial design (0), or block design (1). A study was defined as a block design if multiple trials in the same condition were used consecutively, a tactic often used to maximize signal, and trial design for all others. The number of trials/blocks was a continuous variable, defined as the number of trials and blocks that were separated from other segments by a rest period or a fixation period. The length of trials/blocks was a continuous variable, defined as the duration of each experimental trial/block (i.e., not including baseline or ISI periods). Presence of stimuli was a binary variable, defined based on whether the study was conducted with stimuli (1) or without stimuli (0). Finally, the type of stimuli was a categorical variable, defined as audio (0), visual (1), or audiovisual (2); Here, we only coded for the stimuli specifically designed in the task and did not code for non-task stimuli that may have been present (e.g., attention-grabbers, layout of the lab, etc.). All variables that did not correspond to exactly one of these options were noted and excluded from analyses.

#### 2.4.3 Other Parameters

We also coded for the **final sample size, year of publication**, and **topic of research**. Final sample size was defined as the total number of infants included in each paper’s original analyses. The year of publication was defined as the year in which the publication was accepted. For the topic of research, each experiment was classified as one of the 9 topic categories: Attention, Memory/Learning/Prediction, Auditory/Language, Vision, Multisensory Processing, Social Cognition, Higher Cognition, Resting State, and Other.

### 2.5 Coding

To test our hypotheses outlined in the introduction section, we coded for three types of attrition rates, five Subjects Parameters, six Design Parameters, and three Other Parameters for each experiment. As a result, there were 4,896 possible variables (18 variables x 272 experiments).

Not all variables were reported by all experiments. Variables that were neither reported in the experiment nor calculable based on the information given were noted and excluded from analyses.

For each experiment, two researchers independently coded these variables. Two independent coders agreed on 4,418 of the 4,896 variables, showing an inter-rater reliability of 90.23%. In the case of a disagreement between the two independent coders, a third researcher cast a tie-breaking vote.

### 2.6 Statistical Analysis

All analyses were conducted in R via RStudio (version 4.0.3; R Core Team, 2020). To test our hypotheses about the influence of subject- and design-related variables, we examined whether each parameter of interest independently predicted attrition rate by fitting linear mixed-effects models using the *lme4* package (version 1.1-23; Bates et al., 2015). Models including the parameter of interest as a single fixed effect and publication year as a random intercept were used to predict TAR, IAR, and SAR. While we intuitively assumed that some parameters would be more likely to predict certain types of attrition (e.g., age relating to fussiness such that IAR is affected, or fNIRS cap configuration relating to signal quality such that SAR is affected), we report each parameter’s influence on all types of attrition to paint a full and transparent picture of its effect on data loss.

## 3 Results

### 3.1 Attrition Rates

We examined the overall trends for sample size and attrition rate. Over the 272 experiments, the average final sample size (after attrition) was 23.88 ± 14.47 infants. 220 out of 272 experiments reported enough information to calculate the total attrition rate (TAR). On average, 34.23% of subjects were excluded. Among the 178 experiments that reported reasons for their attrition, 136 experiments reported infant-driven reasons (IAR), and 121 experiments reported signal-driven reasons (SAR). Among studies that reported IAR, 21.50% of the data were excluded due to IAR on average, while among studies that reported SAR, 20.24% were excluded due to SAR on average (Table A; Figure B). Studies that reported IAR or SAR were not significantly different from those that did not in terms of the TAR [*t*(200) = 0.939, *p* = 0.3488]. Our analysis showed that an increasing number of studies are reporting attrition rates; while only 65% of studies reported attrition information between 1998 and 2005, 85% of studies reported attrition since 2005.

**Table A.** Overview of the Attrition Rates

| Type of Attrition | Mean Attrition Rate %<br>*M* (SD) |
| --- | --- |
| Total Attrition Rate (TAR) | 34.23 ($\pm 17.60$) |
| Infant-Driven Attrition Rate (IAR) | 21.50 ($\pm 14.42$) |
| Signal-Driven Attrition Rate (SAR) | 20.24 ($\pm 14.22$) |

**Figure B.**
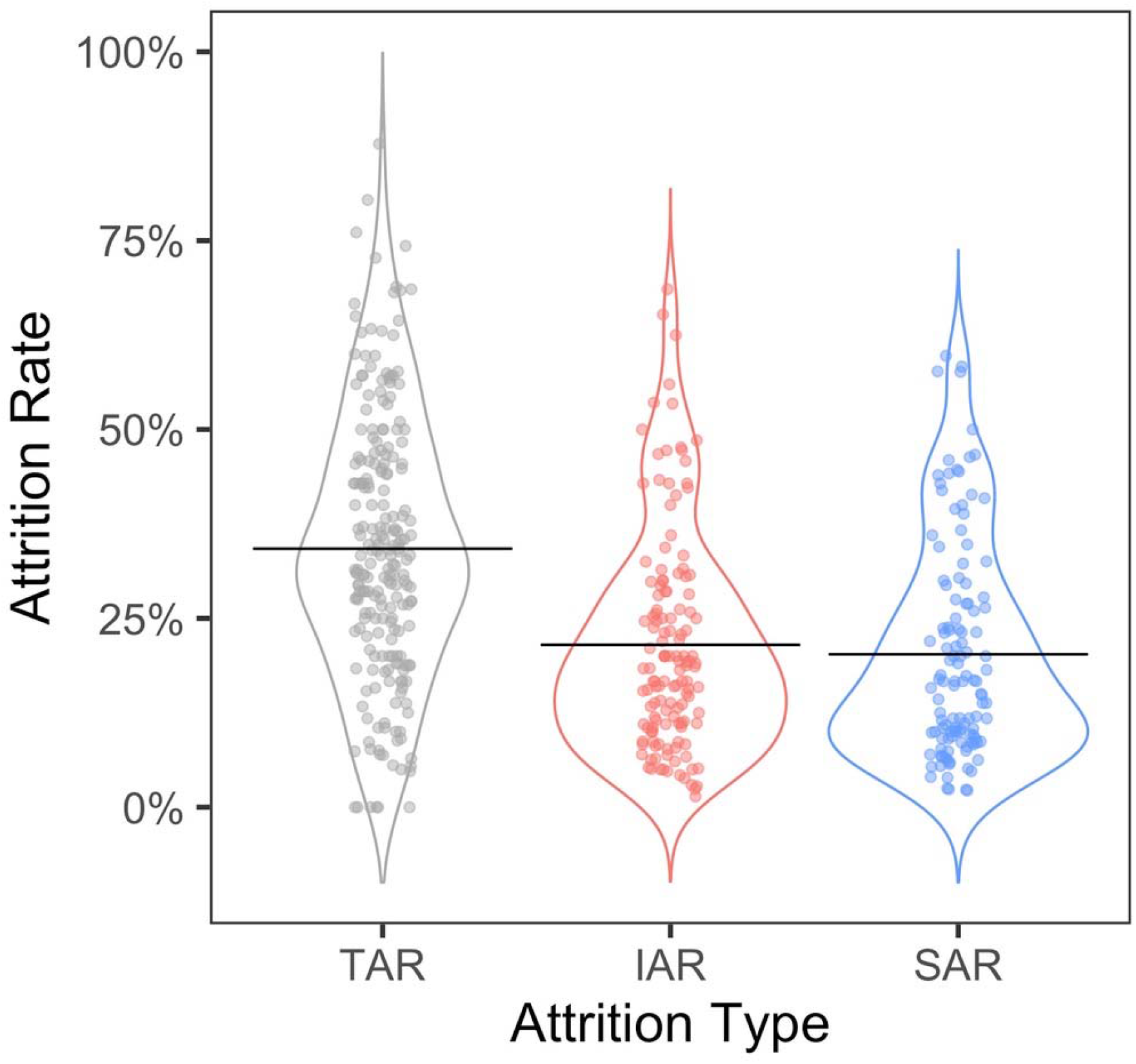
Distribution of attrition types in infant fNIRS research. The distributions of attrition rates reported by all infant fNIRS research to date are depicted for the three types of attrition we analyzed in this report (TAR, IAR, and SAR).

### 3.2 Comparison with attrition rates reported with other research methodologies

One-sample t-tests were conducted to determine if a difference existed between the attrition rates in infant fNIRS and in other methodologies. We compared our TAR/IAR values and the values reported in previously mentioned meta-analyses. Our TAR value of 34.23% was significantly lower than the Stets and colleagues estimate of attrition for infant EEG: 47.3% [t(222) = 14.98, p < 0.001]) suggesting that infant fNIRS studies have less attrition than EEG. Though, Stets et al (2012) did not break down their attrition rates into subtypes (e.g., IAR, SAR) so we are not able to determine which aspect(s) of attrition differ between the modalities. FNIRS attrition rate is significantly higher than the 22.6% attrition rate reported by Slaughter and Suddendorf (2007) for infant looking time studies [(t(222) = 4.89, p < 0.001]). However, as we conceptualized in our approach, fNIRS research likely has multiple sources of attrition including from signal issues as well as infant compliance. To see whether IAR is similar across these methodologies, we compared the IAR we found in fNIRS research (21.50%) to the overall infant-related attrition rate reported by Slaughter and Suddendorf in 2007 (13.7%). We found that there was no significant difference [t(198) = 0.51, p = 0.608)]. This similarity suggests that types or sources of attrition might be similar across methods and literatures.

#### Change in attrition rate over time

The number of infant fNIRS research publications by year has been exponentially increasing since the inception of fNIRS as an infant neuroimaging technique in 1998 (Figure C.1). Considering these trends by research area, the increase was the most pronounced for research in two categories, auditory processing and language and social cognition, but was observed in all explored topics of research (Figure C.2). Given the incredible expansion of the field, we examined changes in the attrition rates as a function of publication year using simple linear regression. We found no significant change in TAR over the years (β = - 0.39, *t* = -1.45, *p* = 0.148, R^2^ = 0.01). However, as hypothesized, there was a significant decrease in SAR over time (β = -1.19, *t* = -4.28, *p* < 0.001, R^2^ = 0.13), with IAR remaining stable (β = - 0.21, *t* = -0.73, *p* = 0.469, R^2^ = 0.00; Figure C.3).

**Figure C.**
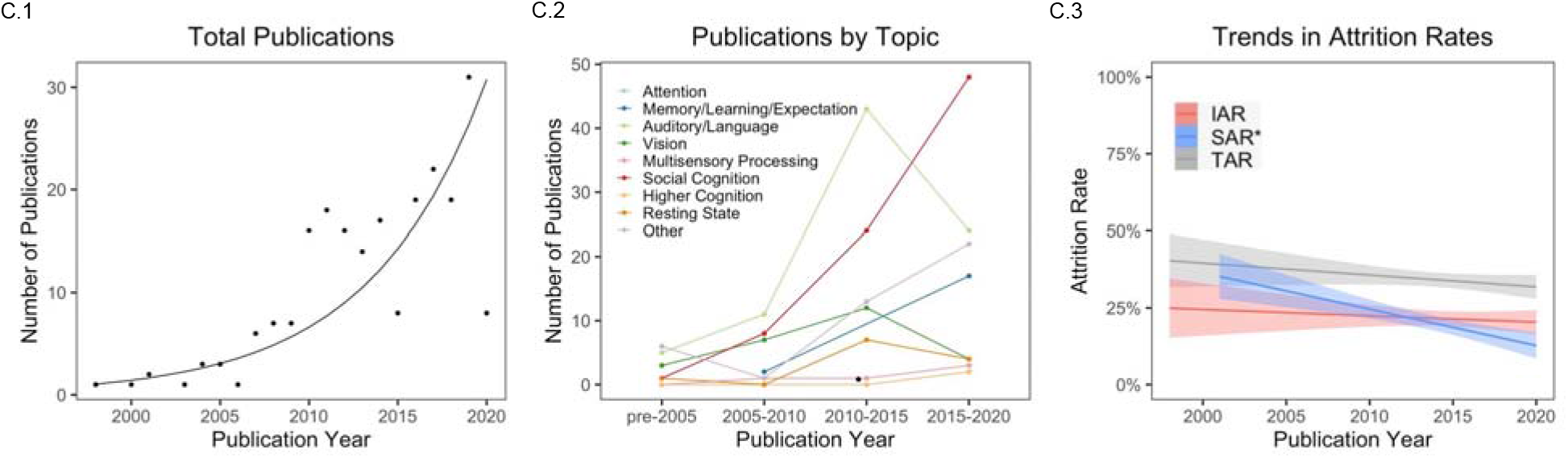
Trends in infant fNIRS research over time. (C.1) Since its inception as an infant neuroimaging technique, fNIRS has been used in an increasing number of infant cognitive neuroscience experiments. (C.2) There has been a general increase in the number of publications on all topics, but the increase was the most pronounced for research on auditory processing/language and social cognition. (C.3) All three types of attrition rates have generally been decreasing over the years, but the only significant decrease was in the attrition rate driven by signal quality (SAR).

We also examined whether final sample size has changed over the history of infant fNIRS studies. The average sample size per experiment significantly increased from *n* = 12.91 in the beginning to *n =* 21.90 in 2010 [t(112) = 2.55, p = 0.030] and has remained steady from *n* = 24.26 to *n* = 24.14 in the last decade ([t(115) = 0.06, p = 0.950].

### 3.3 Subject-Related Parameters

Table B presents a brief summary of the subject-related parameters across all experimental groups (*n* = 272).

**Table B.**
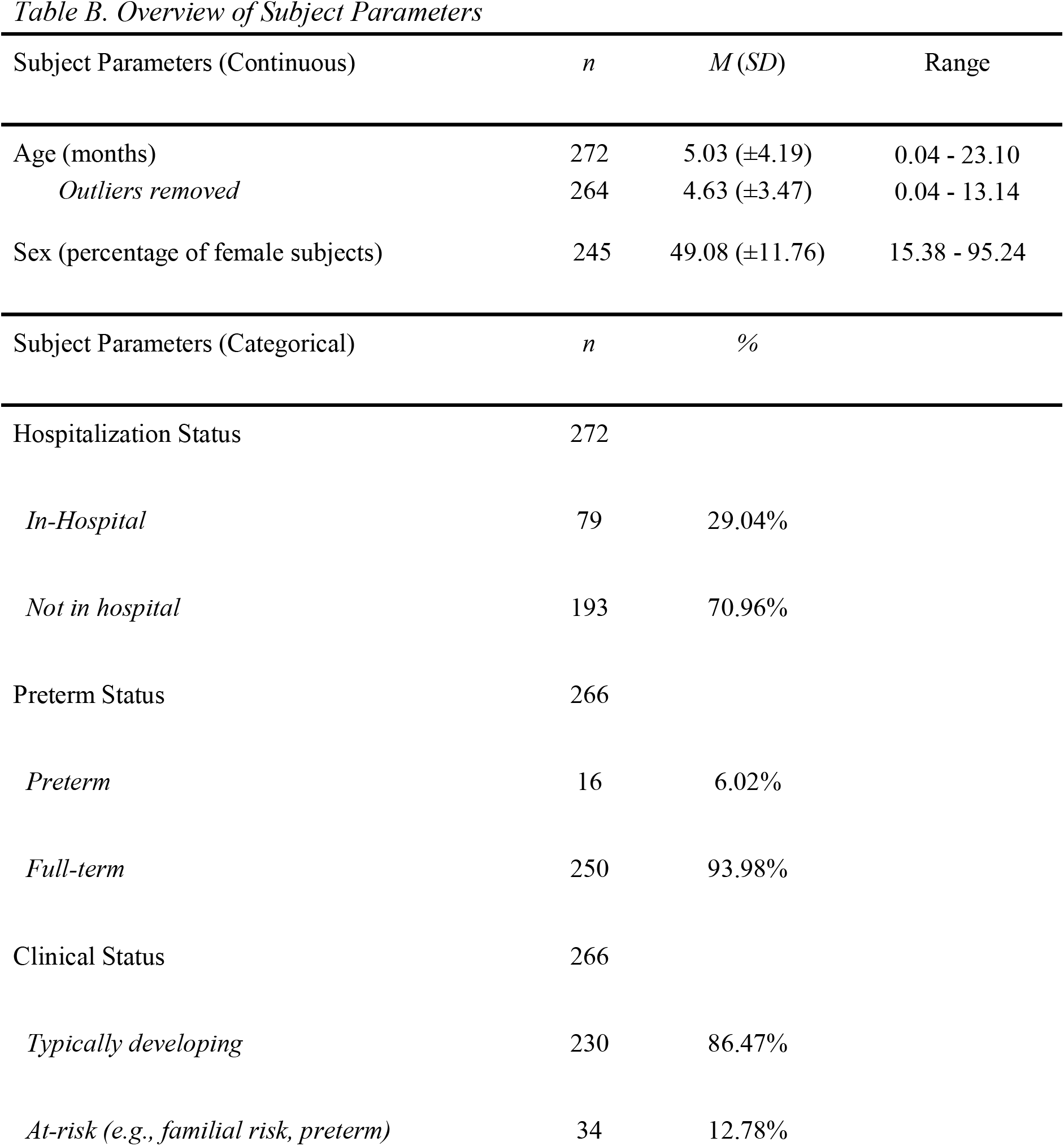
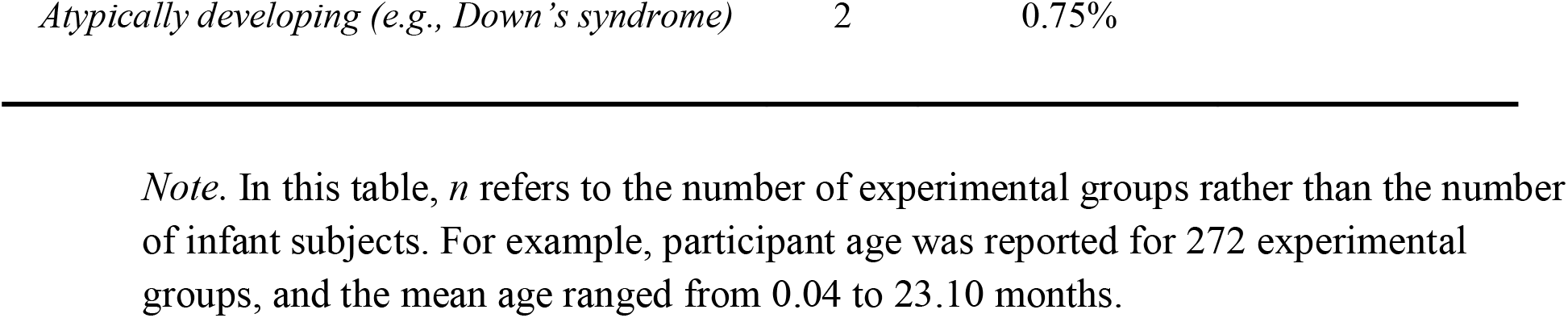
Overview of Subject Parameters

First, we examined the age distribution for all studies (Figure D). Across 272 experiments, 9,114 total infants were recruited, with a mean age of 5.03 ± 4.19 months (range: 0.04-23.10). Age is not normally distributed throughout this range as is found throughout the field of developmental cognitive neuroscience (Azhari et al. 2020). Indeed, seven experiments included infants whose average age was greater than 2 standard deviations away from the overall mean. These outliers (mean age >13.41 months) were retained in the analyses reported below, but we note any instance where the findings diverge for the full sample versus partial sample (excluding outliers).

**Figure D.**
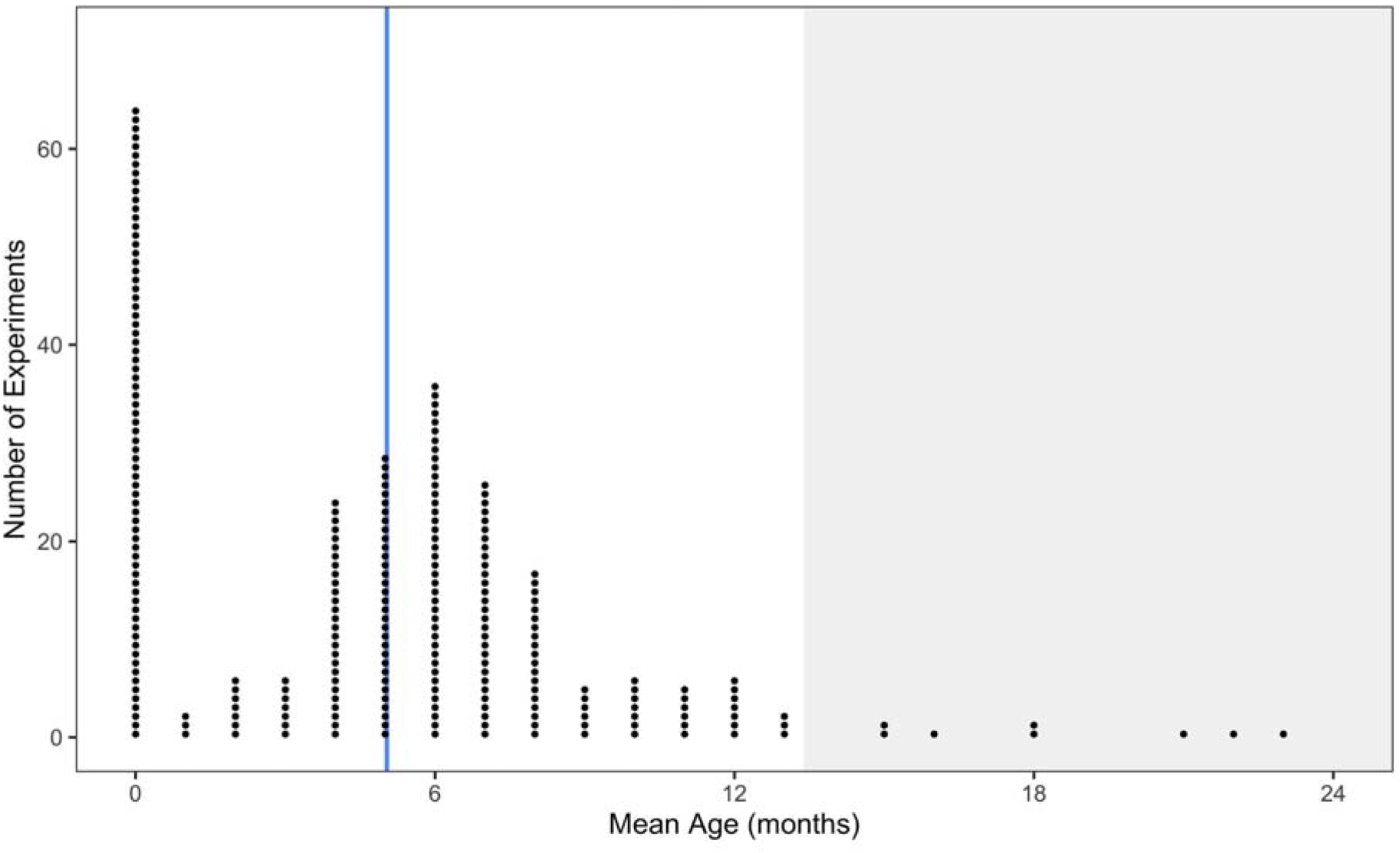
Age distribution for all infant fNIRS experiments. Each point represents a single experiment. The vertical line at 5.03 months reflects the mean age across all experiments. The shaded gray area beginning at 13.41 months denotes the outlier region, defined as >2SD away from the mean age across all experiments.

To test whether participant age influenced attrition rates, we fit a linear mixed-effects model predicting attrition with mean age (in months) as a fixed effect and publication year as a random intercept. Results indicated that age was unrelated to TAR (β = -0.33, *t* = 1.15, *p* = 0.253, R^2^ < 0.01), likely due to the conflicting effects of age on IAR and TAR. Age was a weak but significant positive predictor of IAR (β = 0.76, *t* = 2.70, *p* = 0.008, R^2^ = 0.05; Figure Ea) and weak but significant negative predictor of SAR (β = -0.83, *t* = -2.86, *p* = 0.005, R^2^ = 0.06; Figure Eb). These age effects on IAR (β = 0.87, *t* = 2.40, *p* = 0.018, R^2^ = 0.04) and SAR (β = -0.77, *t* = - 2.04, *p* = 0.044, R^2^ = 0.04) held after removing the oldest outliers from the sample. As a robustness check, we also re-ran these analyses after removing newborns from the sample since there are many differences in the types of experimental tasks used for newborns and older infants (e.g., sleeping vs. awake and behaving), and there is a large concentration of the sample at birth. Only the age effect on SAR remained statistically significant and, in fact, increased in magnitude (β = -1.30, *t* = -3.58, *p* < 0.001, R^2^ = 0.12). Thus, across all three analyses we found that age significantly predicts SAR in the negative direction, such that increased age of infant participants is associated with decreasing attrition due to signal quality issues. The estimates for the effects of age on SAR ranged from β = -0.33 to -1.30.

**Figure E.**
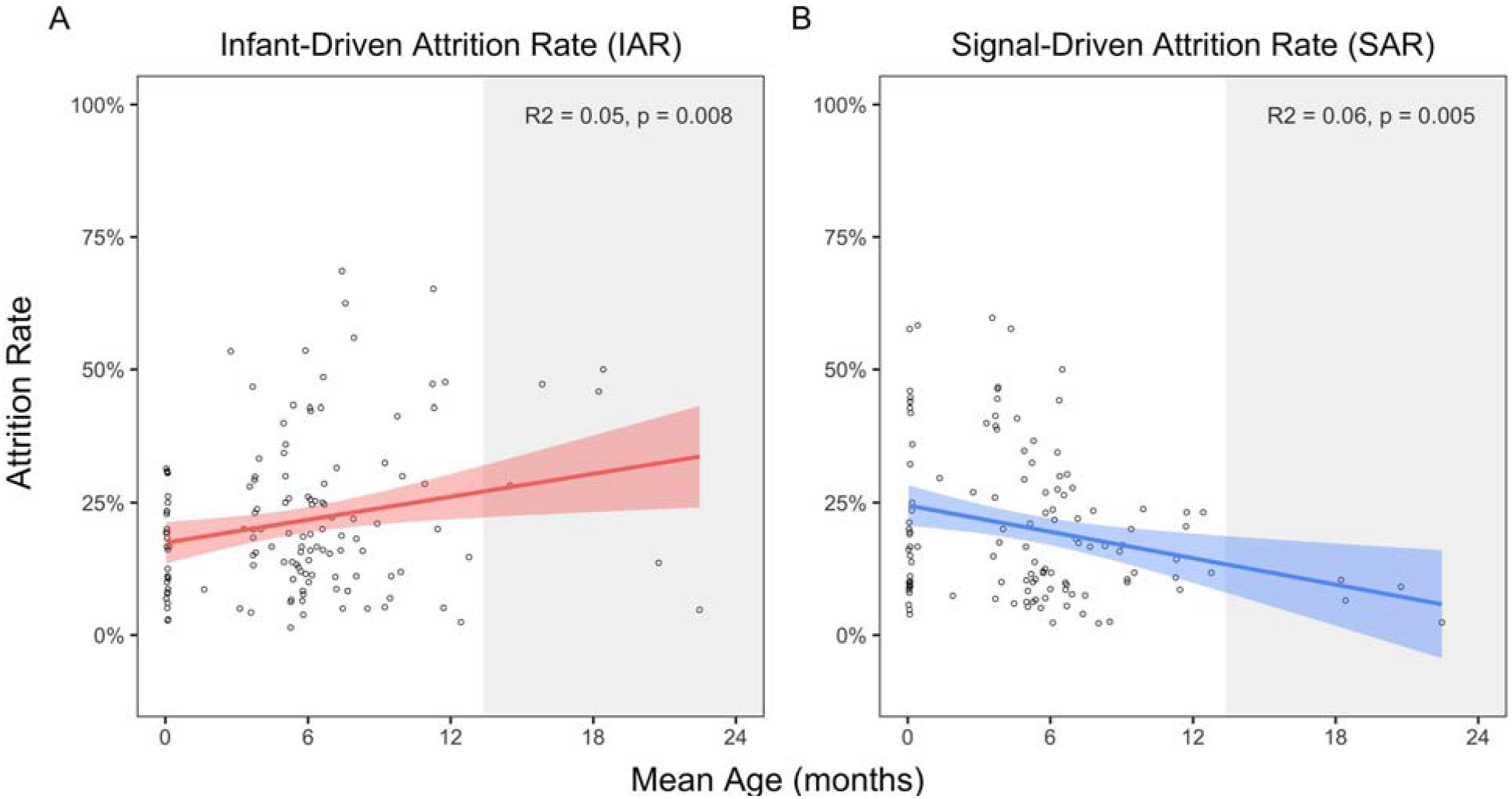
Change in Infant-Driven Attrition Rate (IAR) and Signal-Driven Attrition Rate (SAR) as a function of participant age. (a) Attrition rate due to infant behavior increased with age. (b) Attrition rate due to signal quality decreased with age. The shaded gray area beginning at 13.41 months denotes the outlier region, defined as >2SD away from the mean age across all experiments.

To further explore the effects of age and consider current findings in relation to past studies, we parsed age into categorical bins to determine whether there exists a non-linear influence of age. We first examined whether there were differences in attrition across four age groups (0-2 months, 3-4 months, 5-8 months, 9 months and older; consistent with Cristia et al., 2013) via a one-way ANOVA. Results revealed a significant effect of age on TAR [*F*(3, 216) = 3.33, *p* = 0.021] and SAR [*F*(3, 117) = 6.70, *p* < 0.001], but not IAR. To better understand these significant effects, we performed follow-up pairwise *t*-tests. First, replicating the age effect observed by Cristia et al. (2013), we found that TAR for the 3-to 4-month-old age range was significantly higher than that of the 0-to 2-month group (*p* = 0.003) and the 5-to 8-month group (*p* = 0.019), with only the 0-to 2-month difference remaining significant after Tukey adjustments. For SAR, we similarly found that the 3-to 4-month-old age range experienced higher rates of data loss due to poor signal quality than the 0-to 2-month group (*p* = 0.012), the 5-to 8-month group (*p* < 0.001), and the 9-month and older group (*p* < 0.001), all of which remained significant after Tukey adjustments. Additionally, we found that 0-to 2-month-old age range also experienced higher SAR than the 9-month and older group (*p* = 0.016), though this effect did not hold after correction for multiple comparisons. Together, these findings suggest that studies with younger infants, particularly in the 3-to 4-month age range, may experience higher rates of data loss, particularly due to signal quality issues.

However, this categorization of ages adopted from Cristia et al. (2013) led to uneven group sizes in our sample, with the 3-to 4-month group being the smallest. We therefore re-ran these analyses with updated bin sizes to more systematically sample the 0-to 24-month range (newborns, 1-5, 6-11, 12-17, 18+). With these updated groups, we found no statistically significant categorical differences for any type of attrition via a one-way ANOVA, suggesting that the linear trends reported above could best capture the effects of age.

To test the hypothesis that subjects’ sex affects attrition rate, we calculated the proportion of female subjects from the total sample size for each experiment and tested whether this variable predicted unique variance in attrition rates via mixed-effects regression modeling, with proportion of female subjects as a single fixed effect and publication year as a random intercept. We found that the proportion of female subjects was not a significant predictor for any type of attrition, suggesting that subjects’ sex is generally unrelated to attrition.

We next tested whether there were differences in attrition rate based on the clinical characteristics of subjects. We found that clinical status did not predict significant variance in any type of attrition based on a linear mixed-effects model including clinical status as a single fixed effect and publication year as a random intercept, and we also found no categorical differences in TAR [*t*(218) = 1.12, *p* = 0.265], IAR [*t*(130) = -0.13, *p* = 0.592], or SAR [*t*(115) = 0.99, *p* = 0.323] among typically developing infants vs. at-risk infants. Similarly, we found that preterm status was not a significant predictor of any type of attrition based on a linear mixed-effects model including preterm status a single fixed effect and publication year as a random intercept, and there were also no categorical differences in TAR [*t*(215) = 0.59, *p* = 0.557], IAR [*t*(133) = 0.23, *p* = 0.820], or SAR [*t*(117) = -0.88, *p* = 0.380] for preterm vs. full-term infants. Finally, we also examined the effect of hospitalization status and found that it was a significant negative predictor of TAR (β = -6.12, *t* = -2.14, *p* = 0.033, R^2^ = 0.02) and IAR (β = -6.97, *t* = - 2.41, *p* = 0.017, R^2^ = 0.04) but unrelated to SAR. Specifically, studies with in-hospital infants had an average TAR of 29.58%, and in-lab studies had a significantly higher average TAR of 35.53% [*t*(218) = -2.09, *p* = 0.038]. It is important to note that these results were likely confounded by age differences, with hospitalized infants (*M* = 0.43, *SD* = 1.40 months) tending to be significantly younger than the typical participants of in-lab studies (*M* = 7.02, *SD* = 3.47 months). Indeed, when looking at the subset of studies with infants up to 7.67 months (i.e., the maximum age for in-hospital participants), we found no significant differences in TAR according to testing location based on a Wilcoxon test for non-normally distributed groups (*W* = 2495.5, *p* = 0.075).

### 3.4 Design Parameters

We next examined the impact of the designs of infant fNIRS experiments on the attrition rates (see summary of these variables in Table C).

**Table C.**
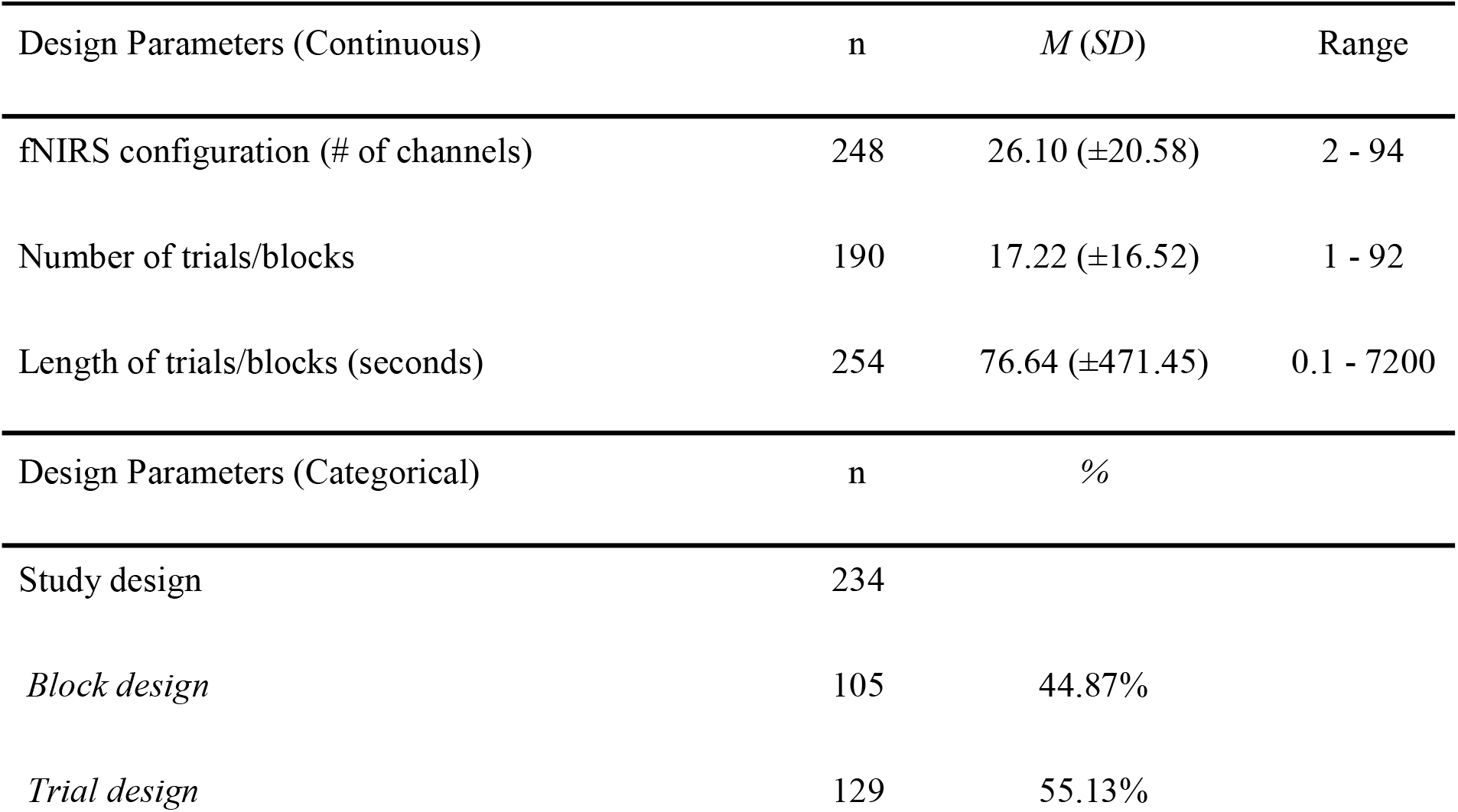
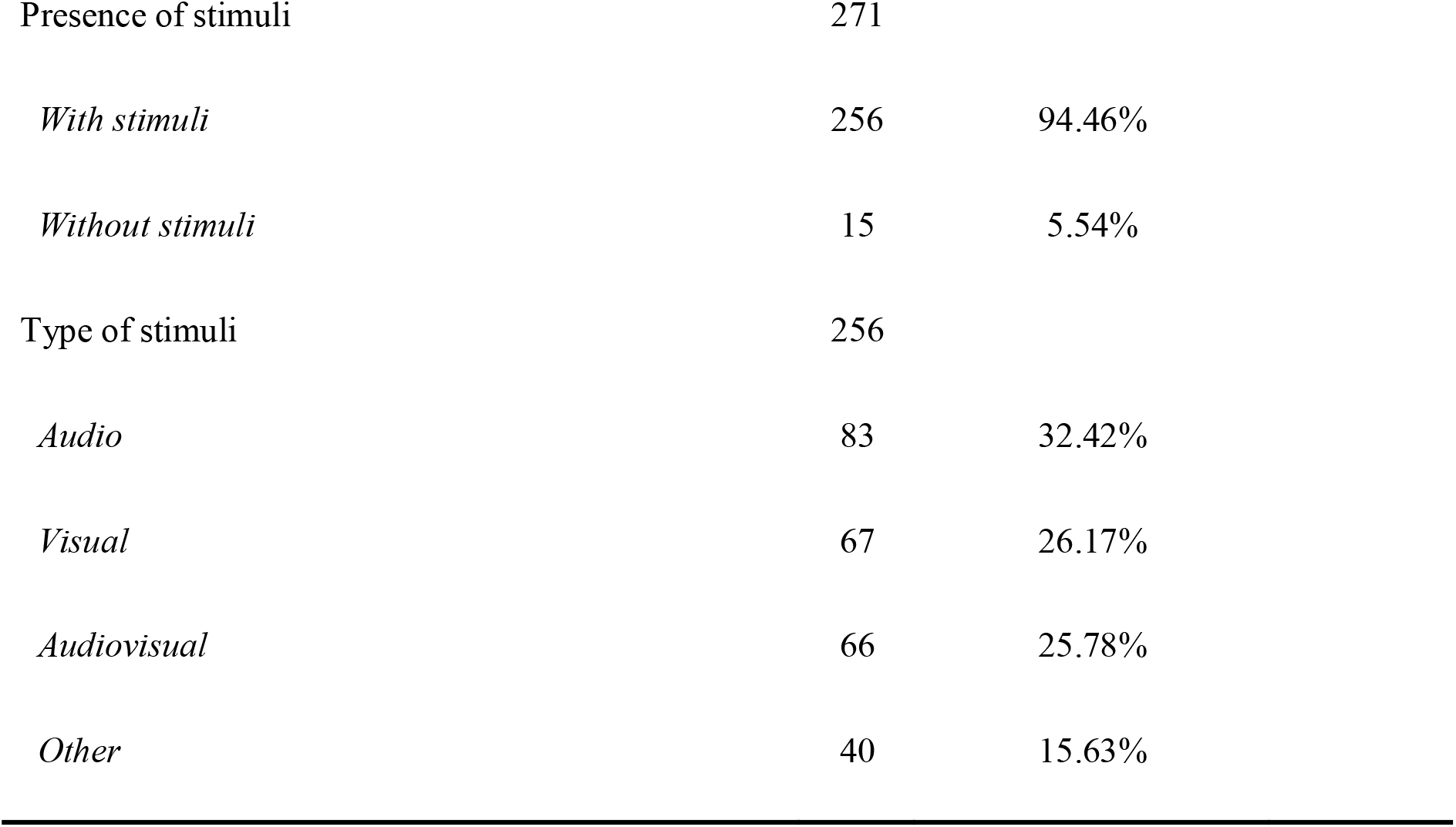
Overview of Design Parameters

| Design Parameters (Continuous) | n | *M* (*SD*) | Range |
| --- | --- | --- | --- |
| fNIRS configuration (# of channels) | 248 | 26.10 ($\pm 20.58$) | 2 - 94 |
| Number of trials/blocks | 190 | 17.22 ($\pm 16.52$) | 1 - 92 |
| Length of trials/blocks (seconds) | 254 | 76.64 ($\pm 471.45$) | 0.1 - 7200 |
| Design Parameters (Categorical) | n | % |  |
| Study design | 234 |  |  |
| *Block design* | 105 | 44.87% |  |
| *Trial design* | 129 | 55.13% |  |

|  |  |  |
| --- | --- | --- |
| Presence of stimuli | 271 |  |
| *With stimuli* | 256 | 94.46% |
| *Without stimuli* | 15 | 5.54% |
| Type of stimuli | 256 |  |
| *Audio* | 83 | 32.42% |
| *Visual* | 67 | 26.17% |
| *Audiovisual* | 66 | 25.78% |
| *Other* | 40 | 15.63% |

First, we examined the effect of the fNIRS cap configuration on attrition. We found that the average number of channels used per experiment increased from 12.56 in 1998 to 23.23 in 2010 [t(110)= 2.17, p = 0.057] but stayed stagnant in the last decade [t(113) = 1.75, p = 0.117]. We ran a linear mixed-effects regression analysis predicting attrition as a function of the number of channels, while controlling for both publication year and participant age. Replicating a previous meta-analysis (Cristia et al., 2013), we found that number of channels was a weak positive predictor of attrition (β = 0.24, *t* = 3.73, *p* < 0.001, R^2^ = 0.06). Interestingly, this was primarily driven by infant behavior rather than signal quality: IAR increased linearly with the number of channels (β = 0.16, *t* = 2.89, *p* = 0.005, R^2^ = 0.05; consistent with Cristia et al., 2013, where the inclusion of more fNIRS optodes was associated with higher attrition rates), while the number of channels did not predict significant variance in SAR (β = 0.10, *t* = 1.53, *p* = 0.129, R^2^ = 0.02).

Next, we tested the effects of the experimental task design on attrition. The duration of a study segment averaged 76.75 seconds (range: 0.1 to 7200 secs, n = 251). We found that study design (block vs. trial) was a positive predictor of TAR (β = 6.59, *t* = 2.71, *p* = 0.007, R^2^ = 0.04) and a trending positive predictor of IAR (β = 4.94, *t* = 1.97, *p* = 0.0051, R^2^ = 0.03) but not SAR. Specifically, studies using a block design had an average TAR of 30.54%, and trial designs had a significantly higher average TAR of 37.14% [*t*(195) = -2.71, *p* = 0.007; Figure 6]. In terms of variance accounted for, this is one of the strongest effects observed. Our findings are in contrast to Cristia et al. (2013) which found that presentation of repeated stimuli resulted in higher attrition rates. We demonstrate that block designs, which by definition are repeating stimuli, significantly lowered the overall attrition rate.

**Figure 6.**
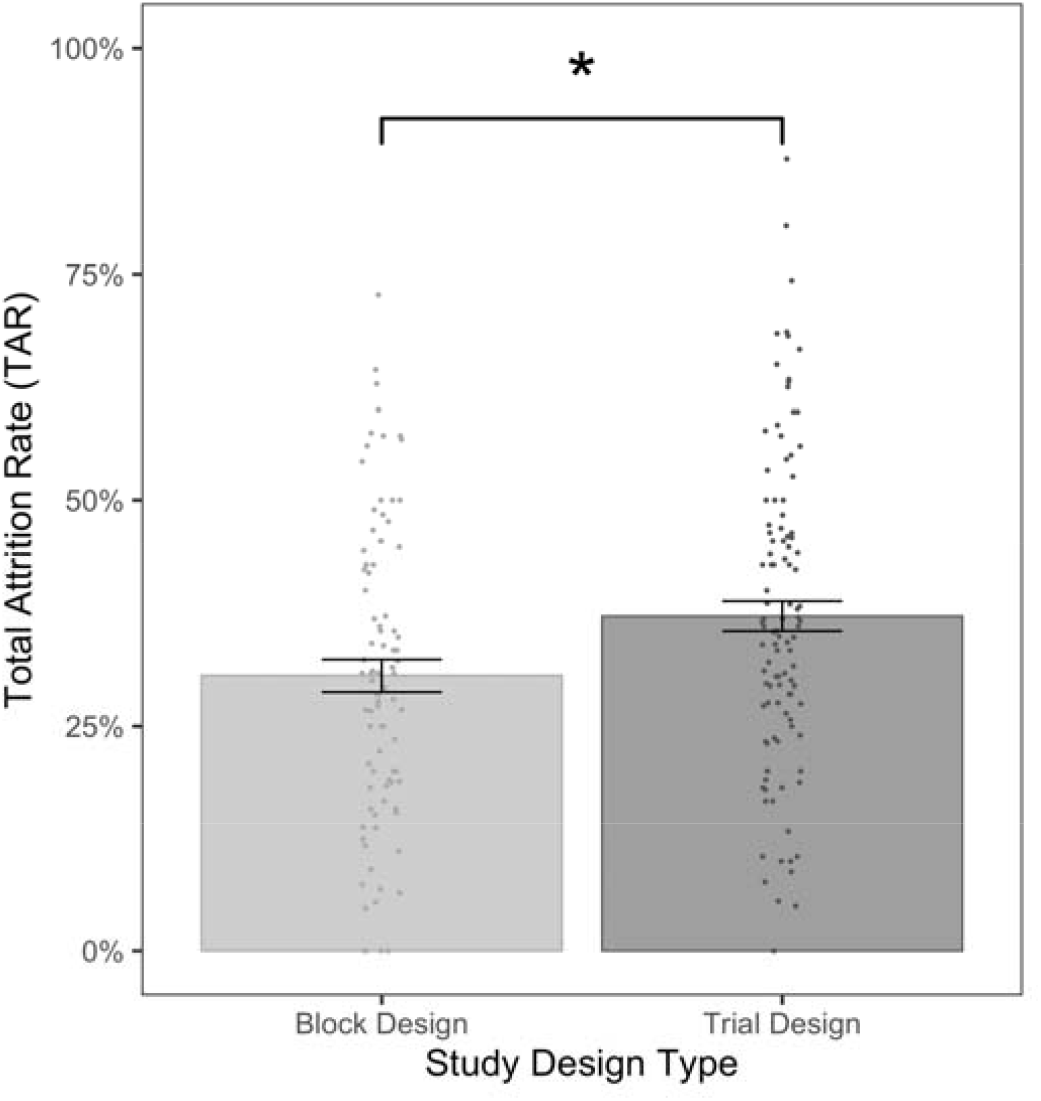
Effect of Study Design Type on Attrition Rate. Studies using a trial design experienced higher attrition rates compared to those using a block design. Points reflect individual TAR means for each experiment, with error bars indicating standard error of the overall mean, * denotes a statistically significant difference.

Next, we examined the effect of the number of study segments (i.e., total number of blocks or trials) on attrition. We fit a linear mixed-effects model predicting attrition, with the number of study segments included as a single fixed effect and publication year as a random effect. Results indicated that number of study segments was a significant negative predictor of both TAR (β = - 0.21, *t* = -2.36, *p* = 0.030, R^2^ = 0.04) and IAR (β = -0.18, *t* = -2.26, *p* = 0.026, R^2^ = 0.05) but not SAR. The amount of variance explained by this parameter is low, and the effect on IAR did not remain statistically significant after removing the oldest age outliers. Given that we found significant differences in TAR and IAR between studies with block vs. trial designs, we also examined the effect of the number of study segments separately for these groups. We found that the effect on TAR held for trial designs (β = -0.23, *t* = -2.18, *p* = 0.033, R^2^ = 0.08) but not block designs, and the effect on IAR was not significant for block or trial designs. Thus, we do not present conclusive evidence that the number of study segments impacts attrition rate.

Given the characteristics of the data, there was not enough variation in this parameter to analyze the effect of study segment duration on attrition. Specifically, the distribution of study segment durations across experiments was highly right-skewed, with only 10 total experiments featuring durations greater than 60 seconds, meaning that outliers would have heavily influenced any observed effects and excluding these outliers would have left little to no variation to examine.

Finally, we examined the effects of experimental stimuli on attrition rates. We found that the presence of stimuli was not a significant predictor of TAR or SAR but was a significant negative predictor of IAR (β = -21.55, *t* = -3.03, *p* = 0.003, R^2^ = 0.06). Specifically, studies including any type of experimental stimulus had an average IAR of 20.87%, and studies not featuring stimuli experienced a higher average IAR of 42.42%. This difference was only trending towards statistical significance based on a Wilcoxon test between non-normally distributed groups (*W* = 405, *p* = 0.070), possibly because of the small sample of studies with no stimulus. However, similar to the trial vs. block designs, this is one of the largest differences in attrition rate, numerically, reported in this meta-analysis.

Then, we compared attrition rates for the three most common types of experimental stimuli (auditory, visual, and audiovisual) using a one-way ANOVA. We found no differences in TAR or SAR according to the type of experimental stimulus but a trending difference in IAR [*F*(2, 122) = 2.92, *p* = 0.058, ηp2 = 0.05]. A follow-up pairwise analysis revealed that studies using audiovisual stimuli experienced marginally greater IAR than studies using only auditory stimuli (*p* = 0.017, adjusted *p* = 0.051; 23.53% vs. 16.67%, respectively). This effect may be at least partially explained by differences in the typical study designs employed for experiments featuring audiovisual vs. only auditory stimuli. Specifically, studies with audiovisual stimuli were more likely to use trial designs, which are associated with higher rates of attrition (68.75% trial designs for audiovisual studies, relative to 24.53% for auditory-only studies). Further research would be needed to tease apart these two possible inter-related factors (study design and stimulus type). There were no significant differences between auditory and visual stimuli (*p* = 0.249; 16.67% vs. 20.40%, respectively) or visual and audiovisual stimuli (*p* = 0.304). Additionally, we found no differences in IAR regardless of whether visual stimuli were static or dynamic (*W* = 985, *p* = 0.328).

## 4 Discussion

A detailed understanding of attrition is essential for any field. It allows researchers to characterize external validity, to plan sample sizes in pre-registered studies, and to reduce attrition by understanding the relevant experimental factors. In this meta-analysis, we built on past work to present a detailed examination of attrition rates across 272 experiments in all 182 infant fNIRS studies published before April 9, 2020. We had two main goals: One was to present a clear characterization of the state of the field concerning attrition and to provide benchmarks upon which future studies could be compared; the second was to examine what factors influence attrition to provide insight for researchers when designing future studies. To this end, we have launched an attrition rate calculator (**LINK**) that will aid researchers in using this meta-analysis to plan future studies, and the database of all of this information is publicly available (https://github.com/soribaek/Attrition-Rate-in-Infant-fNIRS-Research).

Overall, we found that infant fNIRS studies experienced 34.23% attrition. Compared to corresponding adult research (Lloyd-Fox et al., 2010), infant fNIRS research reported greater attrition. Compared to infant EEG (the most comparable methodology), fNIRS research reported significantly lower attrition rate. This difference between fNIRS and EEG is surprising given many similarities between the methodologies. However, the sources of these differences remain unclear given that the meta-analysis of infant EEG research did not consider sub-types of attrition (e.g., SAR). Compared to behavioral infant research, fNIRS research reported greater attrition, but considering the common type of attrition across these methodologies (IAR in fNIRS research vs. infant non-compliance in behavioral research) revealed no significant difference. This similarity in IAR across these methodologies suggests that certain sources of attrition might be constant across modalities, although the current meta-analyses in EEG and infant eye-tracking are unfortunately not sufficiently detailed to allow these comparisons.

We found a general downward trend in attrition over the years, and that this decrease was attributable to fNIRS signal factors (SAR, as opposed to IAR). This trend likely stems from improvements in signal processing (Cope et al., 1988; Cui et al., 2010; Virtanen et al., 2011) and the development of advanced toolboxes (Koh et al., 2007; Hassanpour et al., 2014; Huppert et al., 2009; Fantini, 2014; Tsuzuki & Dan, 2014) and shows promise for continued improvements in the future. Similar findings were observed in a meta-analysis on infant EEG (Stets et al., 2012), where older articles reported higher attrition rates, but this effect could not be attributed to any particular variable because of the limitations of the meta-analysis. In contrast, Cristia et al (2013) found no change in attrition with years; This difference may be because Cristia et al (2013) did not separate sources of attrition and our study adds 8 more years of studies which is a substantial proportion of studies in this young field.

We examined subject characteristics that could affect attrition rates. As hypothesized, we found that age of participants predicted overall greater attrition. However, when considering sub-types of attrition the picture is more complex. Age was a positive predictor of IAR (i.e., older infants are more likely to be fussy), but a negative predictor of SAR. This negative relationship could be due to the fNIRS cap fitting better on older infants with larger heads or older infants being comfortable with the additional weight of the fNIRS hardware. Overall, our investigation into the relationship between age and attrition rate suggests that there is a compromise between the infant-driven and signal-driven attrition rate.

Our age-related findings are in line with previous work that has implicated subject age as a key predictor of attrition rate in neuroimaging research. Cristia and colleagues (2013) found that older infants show lower overall attrition rates than younger infants, and Slaughter and Suddendorf (2007) found a negative correlation between attrition caused by reasons other than fussiness and the age of the infants. We replicated both of these findings. These findings indicate that research can be improved by tackling different causes of attrition depending on the target age. For example, scientists conducting research on older infants can focus on helping them be less fussy (e.g., by focusing on factors presented in section 5). Meanwhile, scientists conducting research on younger infants can focus on improving the signal quality of the data (e.g., by focusing on cap fit and related factors).

Inconsistent with our hypotheses, we found that clinical characteristics were not systematically related to attrition rates (consistent with Stets et al., 2012). This null findings is in contrast to prior research which found that clinical characteristics greatly influenced how likely adult subjects were to cooperate in an experimental study (Chang et al., 2009; Issakidis & Andrews, 2004; Andreescu et al., 2008). This discrepancy shows that there are stark differences in the predictors of attrition between infant and adult subject populations, and further highlight that scientists need to explore age-conscious methods to reduce attrition rates.

Finally, we determined which aspects of infant fNIRS research influences attrition. Replicating a key finding from Cristia et al (2013), we found that number of fNIRS channels positively predicted infant-driven attrition rate. This relationship is likely because the increased weight and size of the fNIRS cap (Lloyd-Fox et al., 2010) increases in infant fussiness. Understanding this relationship is particularly informative since we see a trend of increasing numbers of optodes over time in our field (consistent with Cristia et al., 2013). Although the development of newer fNIRS systems with more channels allows for better coverage of the cortex and the ability to obtain more fine-grained spatial information within regions of interest, it is important to recognize the trade-off between increased signal potential and increased weight or size of the fNIRS cap, which may lead to more participant movement or fussiness (Cristia et al., 2013; Lloyd-Fox et al., 2010; Watanabe et al., 2008).

Interestingly, stimulus design had the largest numerical impact on attrition. We found that studies using block designs experienced significantly lower attrition rates than studies using trial designs. This finding suggests that using a block design, when possible, is likely to reduce data loss. This finding may appear to contradict established findings, which tend to implicate longer study segments as a predictor of *higher* attrition due to infant behavior. For instance, longer studies lead to greater subject dropout across a wide array of data collection techniques (e.g., surveys: Hoerger, 2010; eye-tracking: Hessels et al., 2015) and long, repetitive experimental blocks may be linked to subject restlessness in fNIRS (Cristia et al., 2013). However, our findings are not in opposition to those results. First, the finding regarding block designs does not refer to the length of the overall experiment but instead the duration of stimulation between periods of rest. Thus, viewing a block of stimuli (mean duration = 71.76 seconds) is not equivalent to participating in a long study. Second, these findings point to a likely to be a non-linear relationship between the length of stimulus presentation and engagement with infants, reflective of both the classic Hunter and Ames model (Hunter & Ames, 1988) and Goldilocks effect (Kidd et al., 2012).

The presence and type of stimuli also substantially influenced attrition rate. Studies which presented stimuli experienced half as much IAR compared to studies which presented no stimuli (21.94% vs. 42.41% attrition). Studies using audiovisual stimuli experienced marginally greater IAR than studies using only auditory stimuli. These results are in line with meta-analytic findings from infant EEG (Stets et al., 2012), which reported that audiovisual stimuli resulted in a 19.9% increase in attrition compared to auditory stimuli. This finding is puzzling, given that audiovisual stimuli are widely believed to be more attention-grabbing than visual-only or audio-only stimuli (e.g., Stets et al., 2012; Brannon et al., 2008) and thus have been employed to increase subject retention (e.g., Nikkel & Karrer, 1994; Snyder et al., 2002). These counterintuitive results could be reconciled by the fact that studies using only auditory stimuli commonly study sleeping infants, where there is reduced likelihood of data loss due to subject fussiness or movement (Stets et al., 2012). Audio-only studies also commonly use block designs (see previous paragraph) which is also associated with reduced attrition. In addition, it is also possible that there are some differences in the data inclusion criteria depending on the stimuli; for example, the data inclusion criteria for an audiovisual study may require that infants have to watch the audiovisual stimuli, which would be more stringent than the criteria for an audio-only study. Considering these findings highlights that experimental decisions are not made in isolation and often are related to several other factors, all of which might influence attrition rate. More research is needed to ascertain the reason for this counterintuitive result.

This study is presently the first comprehensive meta-analysis on the attrition rates in all infant fNIRS research. The most related study is Cristia and colleagues (2013) which included some analyses on the attrition rate in infant fNIRS research as a part of introducing their online database of infant research. However, their study included only the small sample of studies that were uploaded to the database at the time of publication (76 publications and theses), and their analyses did not focus on attrition rate or outline possible ways to mitigate it. We improved upon these limitations by including all infant fNIRS research published to date identified using a comprehensive search approach. This resulted in a sizable sample of 218 publications. Moreover, we focused our analyses on attrition rate. We also identified various subtypes of attrition and their potential predictors in experimental research. This more comprehensive approach allowed us to gather the trends in research and specifically make conclusions about factors that can moderate the attrition rate.

Despite the potential utility of our findings in future research, our meta-analyses had a number of notable limitations. First, there was variance in the sample size among the studies, which made it more difficult to accurately depict the general trends in the attrition. Although we used the ratio of final sample size versus all recruited subjects for simplicity and consistency, a few studies with small sample sizes may have skewed results. Second, there was a large proportion of studies that did not clearly report their attrition rates. This lack of information obfuscates the true relationship between some of the factors we explored and the attrition rate and reduces the strength of transparency of the field. Among 218 infant fNIRS research publications between 1998 and 2020, 34 (15.5%) did not report sample size; among those that did, only 182 reported enough information for us to calculate the total attrition rate. Furthermore, many experiments did not explain the reason for data exclusion. Thus, only 83% of studies in the field provided even the most basic information needed to calculate attrition rates and many more did not include further information necessary to evaluate key sources and causes of attrition. In Section 5.1, we make six recommendations for infant fNIRS researchers with regards to reporting attrition rates moving forward to enable a more transparent reporting of data selection criteria. These recommendations could be useful for researchers in related fields (infant fMRI, EEG, looking time or other behavioral methods). Additional guidelines on reporting fNIRS results can be found in Cristia et al. (2013).

Given the growing popularity of fNIRS as a neuroimaging tool to study infant neuroscience, understanding factors that decrease the attrition rate in an infant fNIRS research study is helpful to reduce the attrition rate and increase the quality and quantity of infant fNIRS research. We therefore combine the findings of this paper and previous research to suggest six recommendations to minimize the attrition rate in future infant fNIRS research. Understanding attrition rate and its related factors will contribute to greater reproducibility and generalizability of the findings in the field in the future.

## 5 Conclusion

### 5.1 Guidelines to report the attrition rates in infant fNIRS research

1. Report the total number of subjects recruited prior to any data exclusion.
2. Report the infant-driven attrition rate (IAR), calculated as the percentage of excluded subjects due to infant behaviors.
3. Report the subject-driven attrition (SAR), calculated as the percentage of subjects who were excluded due to signal-driven reasons (e.g., poor fit of the fNIRS cap, excessive artifacts).
4. When possible, operationalize the criteria by which infants are excluded due to their signal (e.g., what number of seconds does an infant have to look away in order to be excluded for fussiness; use SD of the fNIRS signal to reflect excessive artifacts).
5. Report the percentage of subjects who were excluded due to technical difficulties (e.g., file corruption, program crash), experimenter error, or other reasons.
6. Report the total attrition rate (TAR), calculated as the percentage of all subjects who were excluded from the final analyses.
7. Report the demographics of the population included in as much detail as is feasible (sex, race/ethnicity, socio-economic status, language exposure, etc) as well as the demographics of the population excluded with the same detail (ideally broken down by source of attrition). This step is essential for determining the external validity of your sample and the ways in which attrition is impacting that.

### 5.2 Guidelines to minimize the attrition rate in infant fNIRS research

1. Develop fNIRS caps that are fitted to infants’ head sizes
2. Limit the number of channels in the fNIRS cap to only those that are necessary
3. Increase comfort for the infant subjects
4. Use block designs rather than trials designs, whenever possible
5. Use stimuli rather than no stimuli
6. Seek to minimize behavioral issues when working with older infants (e.g., > 5 months); seek to minimize signal-related issues when working with younger infants (e.g., < 5 months).

## Appendix A

### Scoping Articles

Altvater-Mackensen, N., & Grossmann, T. (2016). The role of left inferior frontal cortex during audiovisual speech perception in infants. NeuroImage. https://doi.org/10.1016/j.neuroimage.2016.02.061

Scopus: YES

Boldin, A. M., Geiger, R., & Emberson, L. L. (2018). The emergence of top-down, sensory prediction during learning in infancy: A comparison of full-term and preterm infants. Developmental Psychobiology. https://doi.org/10.1002/dev.21624 Scopus: YES

Emberson, L. L., Boldin, A. M., Robertson, C. E., Cannon, G., & Aslin, R. N. (2019). Expectation affects neural repetition suppression in infancy. Developmental Cognitive Neuroscience, 37(100597). https://doi.org/10.1016/j.dcn.2018.11.001

Scopus: YES

Emberson, L. L., Cannon, G., Palmeri, H., Richards, J. E., & Aslin, R. N. (2017). Using fNIRS to examine occipital and temporal responses to stimulus repetition in young infants: Evidence of selective frontal cortex involvement. Developmental Cognitive Neuroscience. https://doi.org/10.1016/j.dcn.2016.11.002

Scopus: YES

Emberson, L. L., Richards, J. E., & Aslin, R. N. (2015). Top-down modulation in the infant brain: Learning-induced expectations rapidly affect the sensory cortex at 6 months. Proceedings of the National Academy of Sciences of the United States of America, 112(31), 9585–9590. https://doi.org/10.1073/pnas.1510343112

Scopus: YES

Hyde, D. C., Boas, D. A., Blair, C., & Carey, S. (2010). Near-infrared spectroscopy shows right parietal specialization for number in pre-verbal infants. NeuroImage. https://doi.org/10.1016/j.neuroimage.2010.06.030

Scopus: YES

Lloyd-Fox, S., Blasi, A., Volein, A., Everdell, N., Elwell, C. E., & Johnson, M. H. (2009). Social perception in infancy: A near infrared spectroscopy study. Child Development. https://doi.org/10.1111/j.1467-8624.2009.01312.x

Scopus: YES

Lloyd-Fox, S., Wu, R., Richards, J. E., Elwell, C. E., & Johnson, M. H. (2015). Cortical activation to action perception is associated with action production abilities in young infants. Cerebral Cortex. https://doi.org/10.1093/cercor/bht207

Scopus: YES

Miguel, H. O., Lisboa, I. C., Gonçalves, O. F., & Sampaio, A. (2019). Brain mechanisms for processing discriminative and affective touch in 7-month-old infants. Developmental Cognitive Neuroscience. https://doi.org/10.1016/j.dcn.2017.10.008

Scopus: YES

Werchan, D. M., & Amso, D. (2020). Top-down knowledge rapidly acquired through abstract rule learning biases subsequent visual attention in 9-month-old infants. Developmental Cognitive Neuroscience. https://doi.org/10.1016/j.dcn.2020.100761

Scopus: In database, not retrieved. No mention of fNIRS/NIRS

## Appendix B

### Database Search

fnirs or functional near infrared spectroscopy or nirs or near infrared spectroscopy baby OR babies OR infant OR infants OR toddler*

Scopus 1,911

PsycINFO 275

PubMed 1,626

Total Records 3,812

Duplicates Removed 1,695

Records for Screening 2,117

Databases

Scopus

Date Searched: 04/03/2020

(TITLE-ABS-KEY (fnirs OR “functional near infrared spectroscopy” OR nirs OR “near infrared spectroscopy”) AND TITLE-ABS-KEY (baby OR babies OR infant OR infants OR infancy OR toddler*))

Results: 1,911

PsycINFO

Date Searched: 04/03/2020

(fnirs or functional near infrared spectroscopy or nirs or near infrared spectroscopy) AND (baby OR babies OR infant OR infants OR infancy OR toddler*) Limiters - English; Publication Type: All Journals, Dissertation Abstract Search modes - Boolean/Phrase

Results: 275

PubMed

Date Searched: 04/03/2020

((((“spectroscopy, near-infrared”[MeSH Terms] OR (“spectroscopy”[All Fields] AND “near-infrared”[All Fields]) OR “near-infrared spectroscopy”[All Fields] OR (“near”[All Fields] AND “infrared”[All Fields] AND “spectroscopy”[All Fields]) OR “near infrared spectroscopy”[All Fields]) OR (functional[All Fields] AND (“spectroscopy, near-infrared”[MeSH Terms] OR (“spectroscopy”[All Fields] AND “near-infrared”[All Fields]) OR “near-infrared spectroscopy”[All Fields] OR (“near”[All Fields] AND “infrared”[All Fields] AND “spectroscopy”[All Fields]) OR “near infrared spectroscopy”[All Fields]))) OR fNIRS[All Fields]) OR NIRS[All Fields]) AND (“baby”[All Fields] OR “babies”[All Fields] OR “infant”[All Fields] OR “infants”[All Fields] OR “infancy”[All Fields] OR “toddler*”[All Fields])

Results: 1,626

Limiters:

Language: English

Document Type: articles, conference papers, dissertations and theses

